# OSPREY 3.0: Open-Source Protein Redesign for You, with Powerful New Features

**DOI:** 10.1101/306324

**Authors:** Mark A. Hallen, Jeffrey W. Martin, Adegoke Ojewole, Jonathan D. Jou, Anna U. Lowegard, Marcel S. Frenkel, Pablo Gainza, Hunter M. Nisonoff, Aditya Mukund, Siyu Wang, Graham T. Holt, David Zhou, Elizabeth Dowd, Bruce R. Donald

## Abstract

We present OSPREY 3.0, a new and greatly improved release of the osprey protein design software. osprey 3.0 features a convenient new Python interface, which greatly improves its ease of use. It is over two orders of magnitude faster than previous versions of osprey when running the same algorithms on the same hardware. Moreover, osprey 3.0 includes several new algorithms, which introduce substantial speedups as well as improved biophysical modeling. It also includes GPU support, which provides an additional speedup of over an order of magnitude. Like previous versions of osprey, osprey 3.0 offers a unique package of advantages over other design software, including provable design algorithms that account for continuous flexibility during design and model conformational entropy. Finally, we show here empirically that osprey 3.0 accurately predicts the effect of mutations on protein-protein binding. osprey 3.0 is available at http://www.cs.duke.edu/donaldlab/osprey.php as free and open-source software.

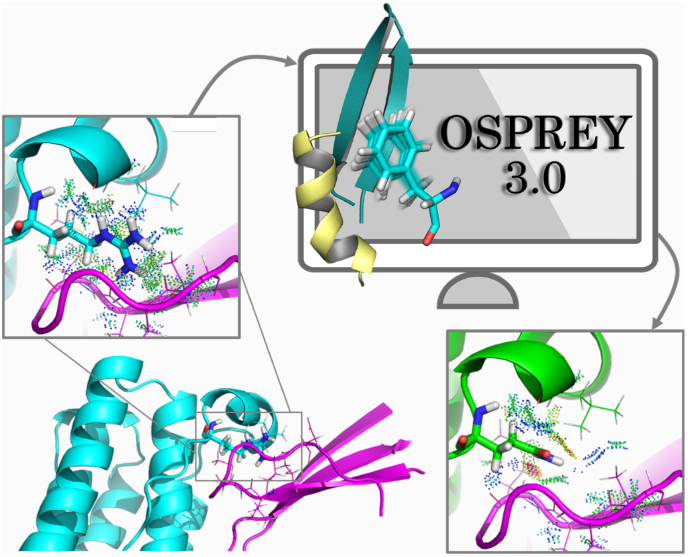

We present the third major release of the OSPREY protein design software, along with comparisons to experimental data that confirm its ability to optimize protein mutants for desired functions. osprey 3.0 has significant efficiency, ease-of-use, and algorithmic improvements over previous versions, including GPU acceleration and a new Python interface.

## INTRODUCTION

For over a decade, the OSPREY software package^1,1–3^ has offered the protein design community a unique combination of continuous flexibility modeling, ensemble modeling, and algorithms with provable guarantees^4,5^. Having begun as a software release for the *K^*^* algorithm ^2,6^, which approximates binding constants using ensemble modeling, it now boasts a wide array of algorithms found in no other software. OSPREY has been used in many designs that were empirically successful—*in vitro*^6–12^ and *in vivo*^7–10^ as well as in non-human primates ^7^. OSPREY’s predictions have been validated by a wide range of experimental methods, including binding assays, enzyme kinetics and activity assays, in cell assays (MICs, fitness) and viral neutralization, *in vivo* studies, and crystal^7,13^ and NMR^9^ structures.

However, as OSPREY grew to include more algorithms and features (Fig. 1), the code became increasingly complicated and difficult to maintain. The growing complexity of the software also hindered its ease-of-use. OSPREY 3.0 represents a complete refactoring of the code, and presents a simpler and more intuitive interface that makes protein redesign much easier than before. The new, developer-friendly code organization also facilitates adding new features to the free and open-source OSPREY project, both by ourselves and by other contributors. We have introduced a convenient Python scripting interface and added support for GPU acceleration of the bulk of the computation, allowing designs to be completed much more quickly and easily than in previous versions of OSPREY. We believe OSPREY 3.0 will be a very useful tool for both developers and users of provably accurate protein design algorithms.

**Figure 1:**
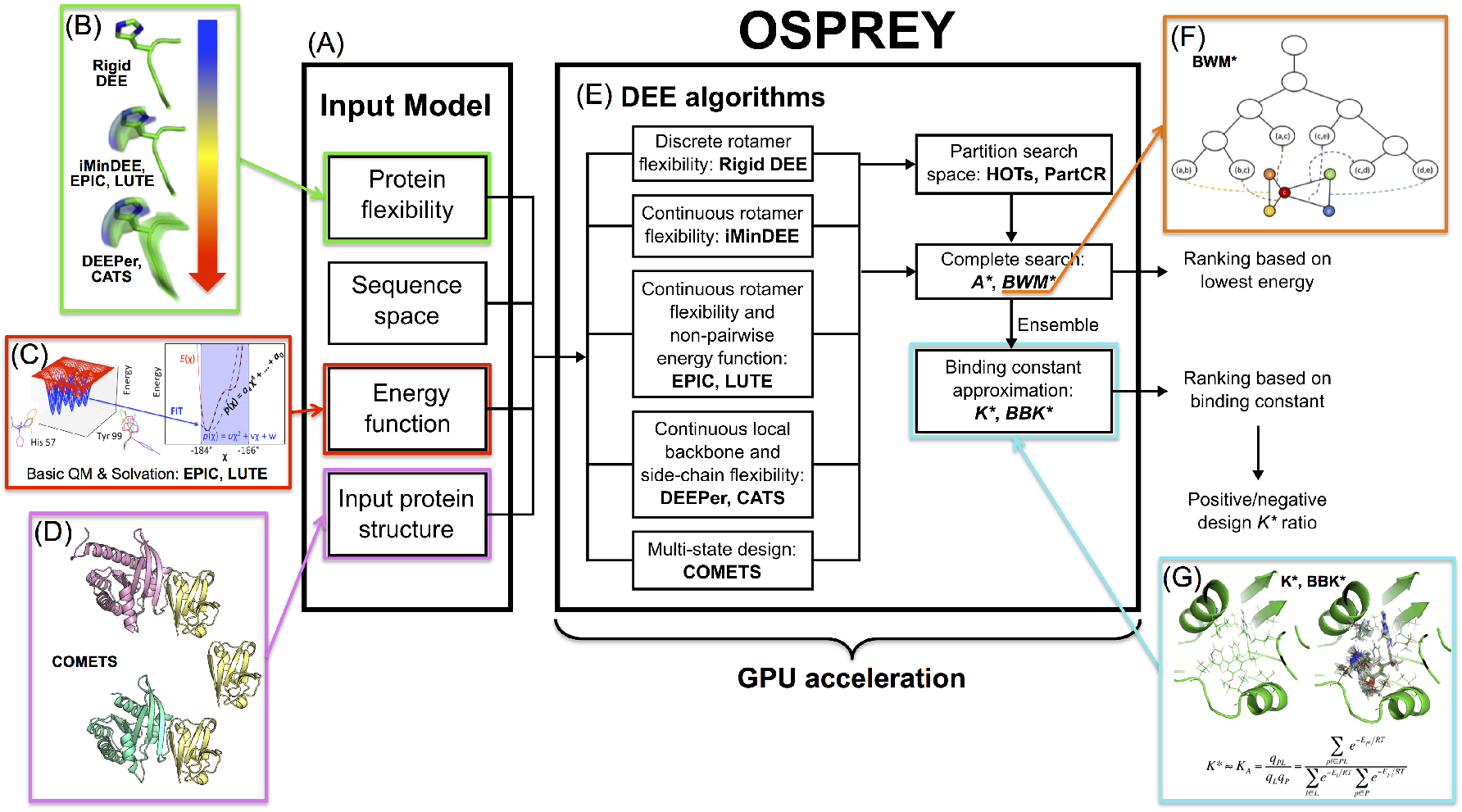
The OSPREY protein redesign suite. (A) The input model includes a 3D structure of the protein to be redesigned, a definition of the sequence space, the allowed protein flexibility (including the rotamer library), and a pairwise energy function. (B) Rigid DEE^14^,^15^, iMinDEE^16^, EPIC^17^, LUTE^18^, DEEPer^19^, and CATS^20^ model different types of protein flexibility. Flexibility ranges in complexity from discrete, rigid rotamers to continuous side chain flexibility to complete flexibility including continuous backbone flexibility. (C) The EPIC ^17^ and LUTE^18^ algorithms also expand energy function capability by allowing for non-pairwise, basic quantum chemistry and Poisson-Boltzmann solvation. (D) COMETS^21^ allows for multi-state design by optimizing sequences and conformations for user-specified bound and unbound states. This is accomplished using multiple input structures. (E) These algorithms are implemented in OSPREY and improved through the use of GPU acceleration. According to the allowed flexibility, OSPREY runs a specific pruning algorithm followed by a highly optimized descendant of the *A^*^* search algorithm^22^. The *A^*^* output generates a ranking based on either the lowest-energy structure of each sequence, or an ensemble of structures computed by the *K*^*^ algorithm. (F) The BWM^*^^23^ algorithm exploits sparse residue interaction graphs and branch decomposition to outperform traditional *A^*^*. (G) The *K^*^*^2,24^ algorithm calculates a *K^*^* score (an approximation of the binding constant, *K_a_*) by provably estimating the partition function for the protein, the ligand, and the protein-ligand complex. The *K^*^* algorithm exploits a thermodynamic ensemble of structures as opposed to a single structure, as illustrated in the panel (PDB ID: 3FQC). *K^*^* can also be used to find sequences that have a high affinity for one ligand (positive design) while having a low affinity for another (negative design) by taking a ratio of *K^*^* scores^10,13^.

### Past successes of osprey

OSPREY has been used for an impressive number of empirically successful designs, ranging from enzyme design to antibody design to prediction of antibiotic resistance mutations. Notably, OSPREY has been successful in many *prospective* experimental studies, i.e., studies in which our designed sequences are tested experimentally, thus validating OSPREY through use in practice rather than simply through a retrospective comparison of OSPREY calculations to previous experimental results. OSPREY is most applicable to problems that can be posed in terms of biophysical state transitions like binding, allowing the *K^*^* algorithm and its variants to predict the optimal sequences based on an estimate of binding free energy computed using Boltzmann-weighted conformational ensembles. Moreover, most protein design problems can be posed in this way, sometimes in terms of binding to more than one ligand. OSPREY is capable of both *positive design*, in which binding of a designed protein to a target is increased, and *negative design*, in which binding to a target is decreased, as well as more complicated design objectives where specific binding to one target and not to another is required.

For example, we have successfully predicted novel resistance mutations to new inhibitors in MRSA (methicillin-resistant *Staphylococcus aureus*) using multistate design (combining negative and positive design). OSPREY does this by searching for sequences that have impaired drug binding compared to wild-type DHFR, but still form the enzyme-substrate complex as usual, allowing catalysis to proceed^10,13^. Our predictions were validated not only biochemically and structurally, but also at an organismal level^13,25,26^. Similarly, we have successfully changed the preferred substrate of an enzyme—the phenylalanine adenylation domain of gramicidin S synthetase—from phenylalanine to leucine by modeling the two enzyme-substrate complexes, and searching for sequences with improved binding to leucine and reduced binding to phenylalanine^6^. The resulting designer enzymes exhibited improved catalysis, and designs changing the specificity from phenlyalanine to several charged amino acids were successful as well^6^. The combination of positive and negative design in OSPREY has also successfully designed mutants of the gp120 surface protein of HIV that bind specifically to particular classes of antibodies, enabling their use as probes for detecting and isolating those antibodies from human sera^12^.

These multistate design capabilities, long a mainstay of OSPREY, are accelerated by the modules *BBK^*^* (described below) and COMETS (described in Ref. 21). COMETS provably returns the sequence that minimizes any desired linear combination of the energies of multiple protein states, subject to constraints on other linear combinations. Thus, COMETS can target nearly any combination of affinity (to one or multiple ligands), specificity, and stability (for multiple states if needed). COMETS and *BBK^*^* have been integrated into OSPREY 3.0 and accelerated, and they are currently the only provable multi-state design algorithms that run in time sublinear in the size *M* of the sequence space. This can be important, since *M* is exponential in the number of simultaneously mutable residue positions.

Further successes of OSPREY have involved improving positive design, e.g., the interaction of the anti-HIV antibody VRC07 with its antigen, gp120. Using this approach, we collaborated with the NIH Vaccine Research Center to design a broadly neutralizing antibody (VRC07-523LS) against HIV with unprecedented breadth and potency that is now in clinical trials (Clinical Trial Identifier: NCT03015181^7,27^). We also have designed allosteric inhibitors of the leukemia-associated protein-protein interaction between Runx1 and CBF*β*^9^. Similarly, we have used OSPREY to develop peptide inhibitors of CAL, a protein involved in cystic fibrosis^8^. The CBF*β* and CAL inhibitors were successful *in vitro* and *in vivo*^8,9^.

In addition, a number of other research groups have successfully used the OSPREY algorithms and software (by themselves) to perform biomedically important protein designs, *e.g*., to design anti-HIV antibodies that are easier to induce^28^; to design a soluble prefusion closed HIV-1-Env trimer with reduced CD4 affinity and improved immunogenicity^29^; to design a transmembrane Zn^2+^-transporting four-helix bundle^30^; to optimize stability and immunogenicity of therapeutic proteins^31–33^; and to design sequence diversity in a virus panel and predict the epitope specificities of antibody responses to HIV-1 infection^34^.

We believe OSPREY 3.0 will enable an even greater range of successful designs.

## PERFORMANCE ENHANCEMENTS IN osprey 3.0

### Engineering improvements yield large single-threaded speedups

OSPREY 3.0’s code has been heavily optimized to improve single-threaded performance relative to the previous version, OSPREY 2.2^21^. Two main areas have received the most attention and the most improvement in performance so far: *A^*^* search speed, and conformation minimization speed.

OSPREY uses the *A^*^* search algorithm^15^ to perform its combinatorial search over sequence and conformational space^2,16,19^. The performance of *A^*^* search in OSPREY depends mostly on the size of the conformation space of the design: the time required for search scales strongly with the number of mutable and flexible residues. Search time is also dependent on the speed at which we can evaluate the energy scoring functions on *A^*^* nodes. Optimizations in OSPREY 3.0 have dramatically increased the *A^*^* node scoring speed, mainly by caching the results of expensive computations and reusing them at different nodes. Many intermediate values used by the *A^*^* scoring functions need only be computed once per design. This reduces the cost of node scoring by roughly an order of magnitude. We can also score child nodes differentially against their parent nodes to speed up node scoring. Caching intermediate values during the parent node scoring and using them to simplify child node scoring yields roughly another order of magnitude speedup in *A^*^* node scoring.

OSPREY 3.0 also includes optimizations to improve the performance of forcefield evaluation and conformation minimization. Conformation minimization is typically the bottleneck in OSPREY calculations with continuous flexibility^2,16,19,20^. The code in OSPREY 3.0 that evaluates forcefield energies for a protein conformation has been heavily optimized, although speed gains here over OSPREY 2 are modest (roughly two-fold), since the original code was already well-optimized in this area. Much larger performance increases were gained by caching forcefield parameters and lists of atom pairs between different conformations to be minimized, which yielded roughly a 10-fold increase in speed. OSPREY 3.0 also increases performance by only evaluating forcefield terms involving mutable and/or flexible residues in a design, since interaction energies between other residues will be identical across all sequences and conformations. Since most designs only model a minority of the residues in a protein as flexible, this can be a substantial improvement.

Performance comparisons are shown for 45 protein design test cases in Fig. 2 and Table 1. All these test cases model continuous protein flexibility^2,16,17^, and 18 of them involve provably accurate partition function calculations (see Table 1 and Ref. 17 for details). To summarize, the optimizations to single-threaded performance described above made OSPREY on average 461-fold faster than OSPREY 2.2 across 29 protein design test cases, and allowed OSPREY 3.0 to finish the remaining 16 test cases, which OSPREY 2.2 could not finish within a 17-day time limit. For example, OSPREY 2.2 on a Intel Xeon E5-2640 v4 CPU took 49.5 minutes to run a small (6 continuously flexible residues) benchmark sidechain packing problem involving a 114-residue fragment of PDZ3 domain of PSD-95 protein complexed with a 6-residue peptide ligand (PDB ID: 1TP5). But OSPREY 3.0 finished the same design in 7.0 seconds on the same hardware, which is a 424-fold speedup.

**Figure 2:**
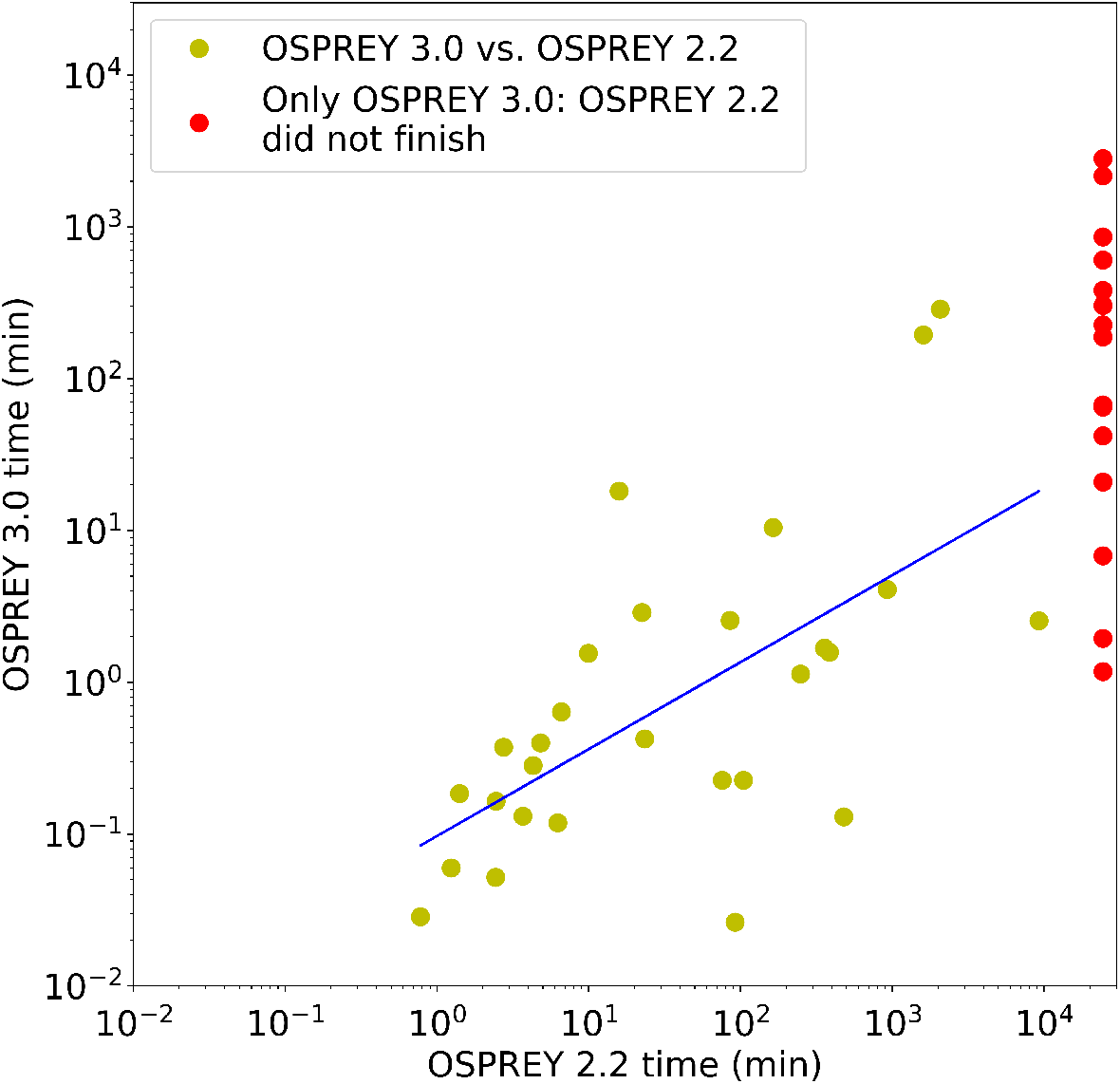
Runtimes of OSPREY 2.2 vs. OSPREY 3.0 for 45 protein design test cases (details shown in Table 1), shown on a log scale. Designs that only finished with OSPREY 3.0 (given a 17-day time limit) are shown on the right in red. All test cases involve continuous flexibility^2,16^ and minimization-aware DEE^16,17^; 18 involve provably accurate partition function calculations (see Table 1 and Ref. 17 for details).

**Table 1:**
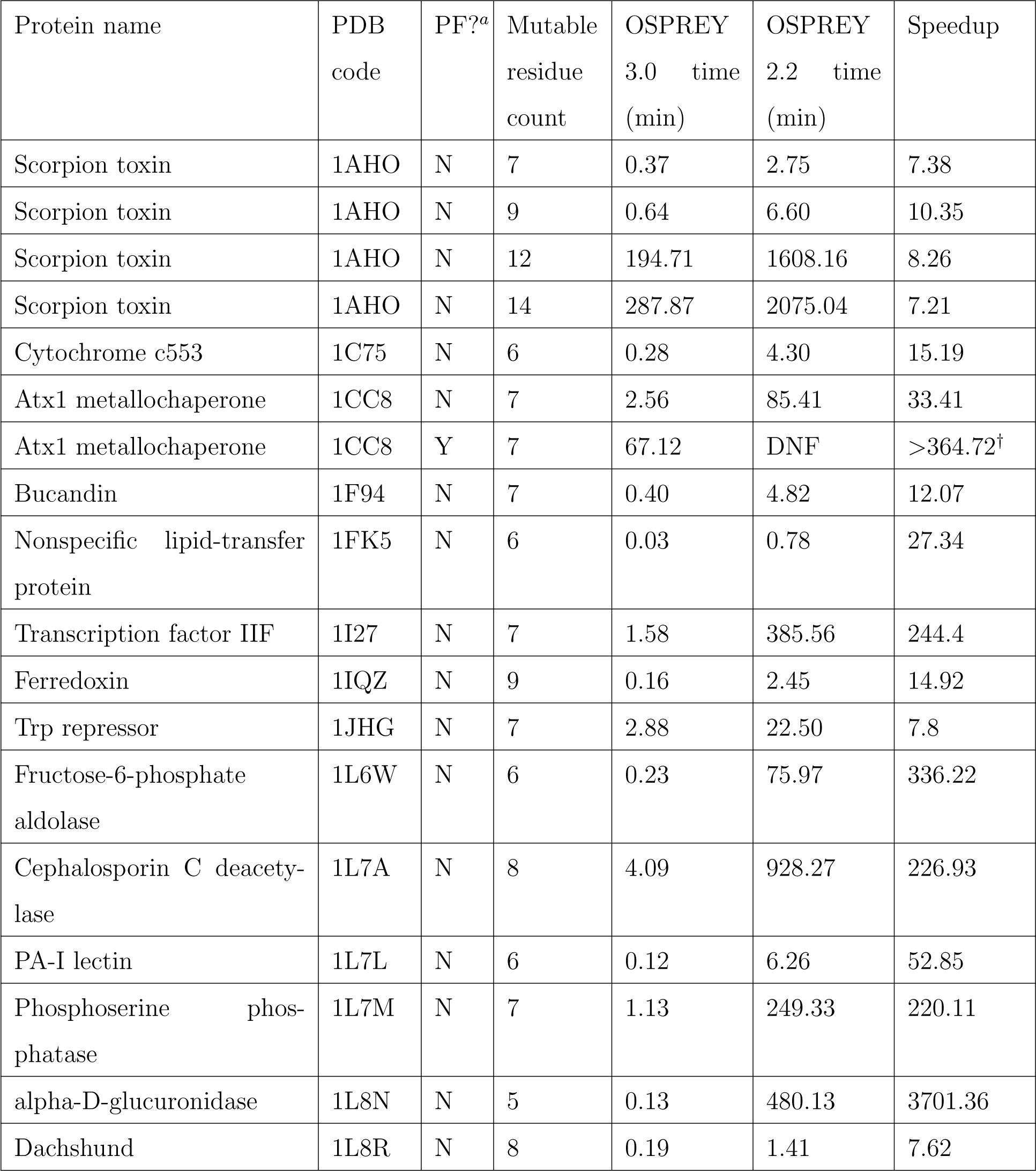
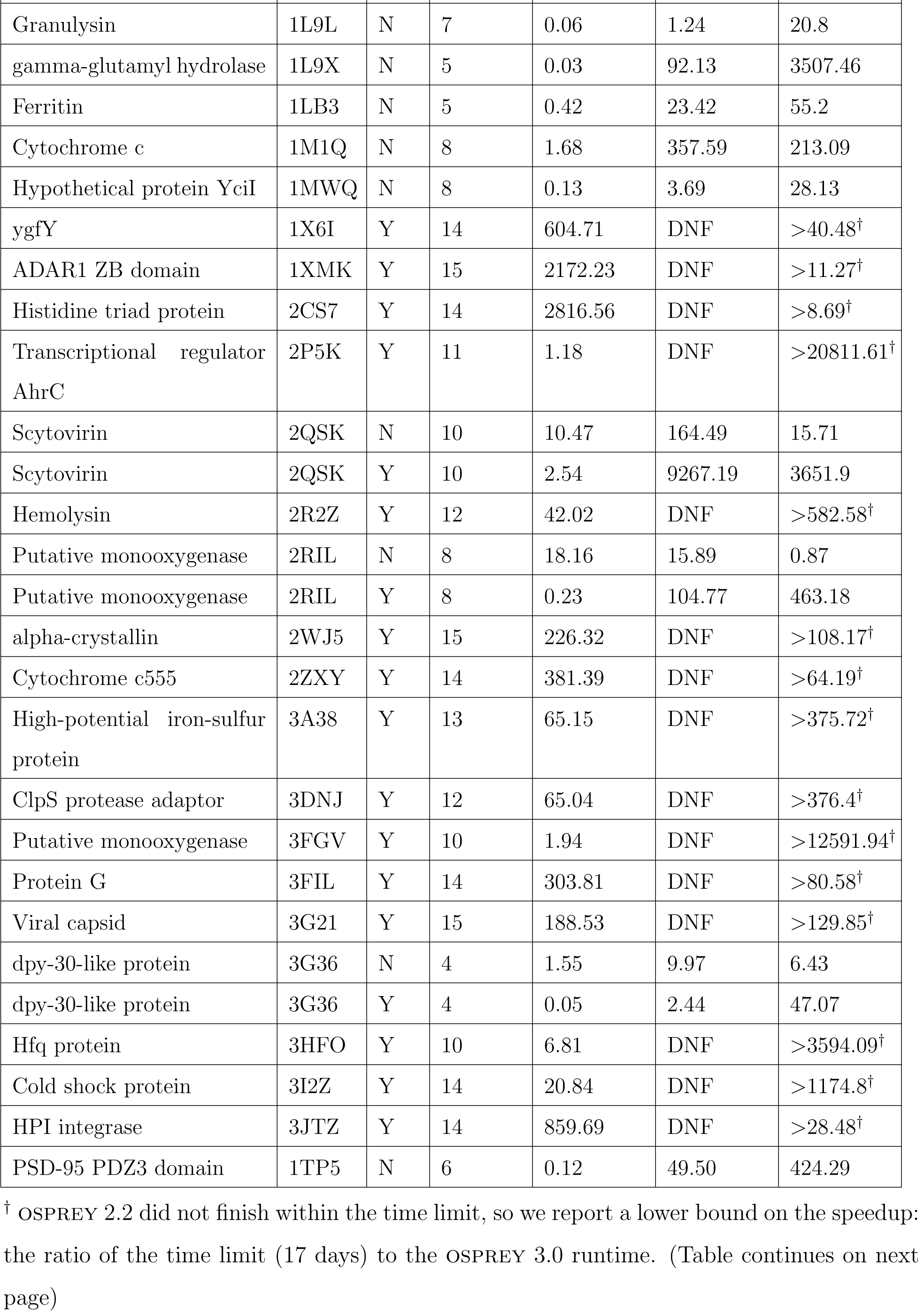
Details of 45 protein design test cases with continuous flexibility run on both OSPREY 2.2 and OSPREY 3.0. Test cases primarily adapted from Ref. 17. ^*a*^ Y indicates a partition function calculation (a subroutine of the *K^*^* algorithm^2,24^), which analyzes a thermodynamic ensemble of conformations; N indicates calculation of the single global minimum-energy conformation (GMEC). DNF: Did not finish.

### GPU acceleration reduces design runtimes

One of the key challenges in protein design is modeling and searching the many continuous conformational degrees of freedom inherent in proteins and other molecules. The sidechain conformations of each amino-acid type are generally found in clusters, known as rotamers^35^, so it is common practice to approximate protein conformational space as discrete by forcing each residue to be in the modal conformation of one of these clusters^14,15^. However, design accuracy is increased significantly when continuous flexibility is taken into account, by allowing the continuous degrees of freedom to move within finite bounds around these modal values^1,16,19,36^. Moreover, this increase in accuracy depends on considering continuous flexibility *during* the conformational search process, rather than simply performing minimization *post hoc* on the top-scoring sequences and conformations output by a discrete search algorithm. Although such a *post hoc* minimization approach would obtain more energetically favorable models of the top sequences, it would still produce the same top sequences as a purely discrete design would, which have been shown to not be truly the top sequences, even if a much finer discrete rotamer subsampling is allowed^1,16^. For example, clashing discrete rotamers can often be converted to favorable conformations by relatively small adjustments in the sidechain conformations^2,16,19,20^. As a result, designs performed with continuous flexibility taken into account *throughout the search* yield significantly different, and more biologically accurate, sequences than the same designs performed using discrete search^1,16,19^.

To address this problem, OSPREY includes several algorithms to design proteins while taking continuous flexibility into account throughout the process of sequence and conformational search^2,16–20^. These algorithms predict optimal protein sequences with provable guarantees of accuracy given a biophysical model that includes continuous flexibility.

This *minimization-aware* design approach requires energy minimization to be performed for a large number of conformations (within the bounds on the continuous degree of freedom that define each conformation). This minimization is a relatively expensive operation, so the bulk of a design’s runtime can be spent on energy minimization of conformations. Therefore, improvements to the speed of energy minimization can have a dramatic impact on OSPREY runtimes.

Much work has been done to optimize OSPREY for execution on CPUs, particularly highly multi-core CPUs and even networked clusters of CPU-powered servers^37,38^. However, modern GPU hardware enables high-performance computation for some specific tasks at a fraction of the cost of large CPU clusters, mainly due to the huge video game industry, which propels innovation in hardware design and drives down costs. The widespread adoption of fast and highly programmable GPUs in the past decade has transformed many areas of computational science, including quantum chemistry^39^, computer vision^40^, and cryptography^41^. In particular, GPUs have been found to produce speedups of approximately an order of magnitude in molecular dynamics simulations^42–44^, which, like OSPREY, must sum huge numbers of forcefield energy terms and can use the GPU to parallelize this computation. GPUs have also been used to accelerate the *A^*^* search step of protein design^45^, albeit without addressing the continuous minimization bottleneck.

Thus, in order to bring the benefit of GPUs to continuously flexible protein design calculations, OSPREY 3.0 includes GPU programs (called *kernels*) built using the CUDA framework^46^ that implement the forcefield calculations and local minimization algorithms used in protein redesign.

We present performance results of these GPU kernels on various hardware platforms in Figure 3. A GPU server housing 4 Nvidia Tesla P100 cards can finish minimizations with about 300,000 atom pairs roughly 110-fold faster than a single thread running on an Intel Xeon E5-2640 v4 CPU. With two Intel Xeon E5-2640 v4 CPUs running at full capacity with multiple threads, the four Nvidia Tesla P100 GPUs finish the same minimizations roughly 8-fold faster. The speedups of GPUs over CPUs scale with the number of atom pairs in the minimization. For minimizations with fewer (about 30,000) atom pairs, even four Nvidia Tesla P100 GPUs cannot outperform two Intel Xeon E5-2640 v4 CPUs. There is significant overhead to transfer each minimization problem from the CPU to the GPU during designs. Even though GPUs can evaluate the minimizations much faster than CPUs, when there are few atom pairs, this transfer overhead dominates the computation time and causes GPUs to perform merely similarly to CPUs, rather than significantly faster.Nevertheless, the bottleneck in protein design is minimizations with many atom pairs, and for these minimizations OSPREY’s speedups on GPUs are on par with the state of the art for GPU speedups of molecular dynamics simulations.

**Figure 3:**
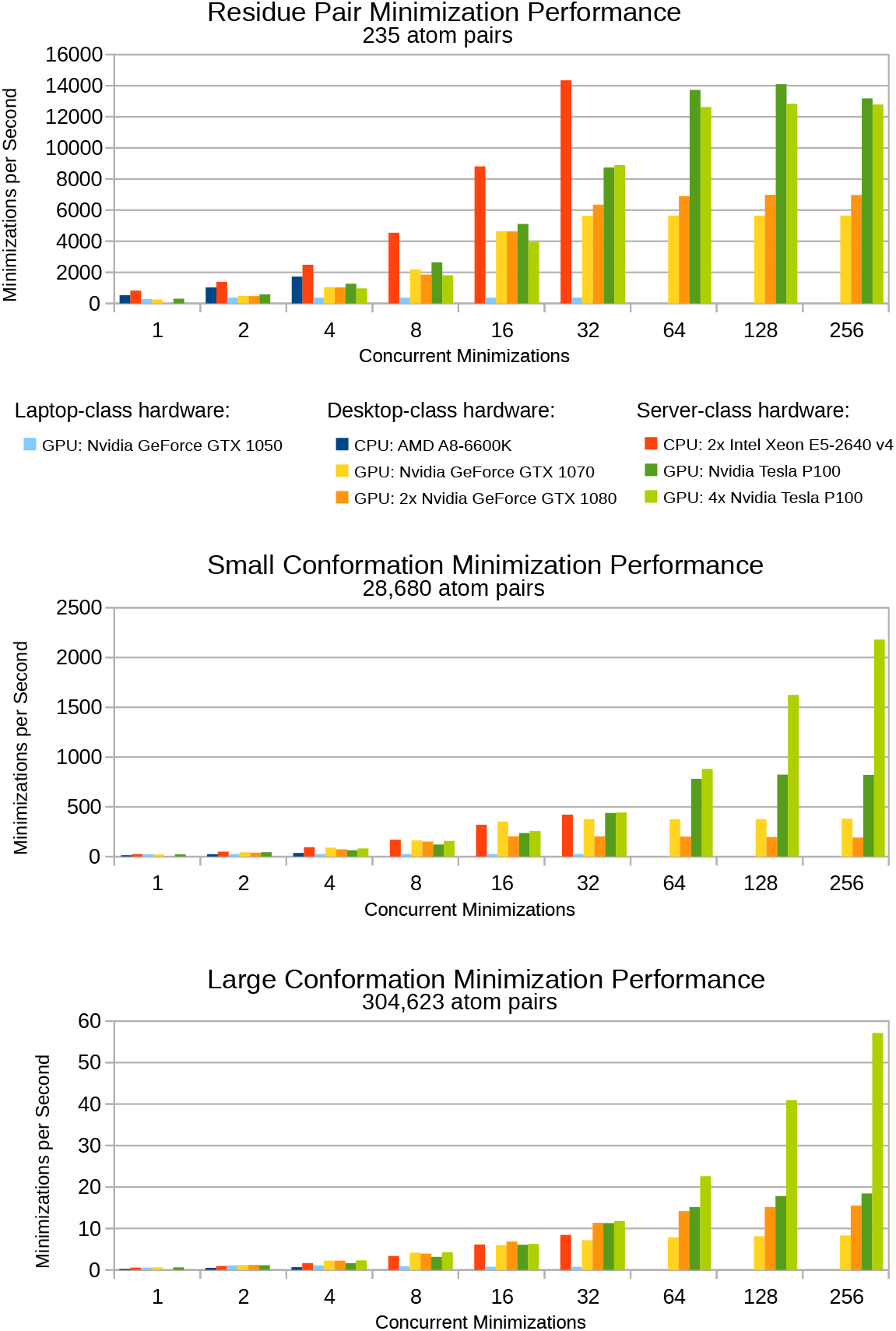
Benchmarks for protein conformation minimization in OSPREY 3.0 for various hardware platforms and for conformations of varying size. From smallest to largest: **(top)** a single residue pair is the smallest multi-body minimization possible, **(middle)** a full protein conformation with a single flexible residue represents a small design, **(bottom)** a full protein conformation with 20 flexible residues represents a large design. For CPU hardware, concurrent minimizations correspond to CPU threads. For GPU hardware, concurrent minimizations correspond to *streams* defined by the CUDA framework. Faster minimization speeds correspond with faster OSPREY runtimes. All minimizations were performed on the Atx1 metallochaperone protein (PDB ID: 1CC8)^47^. Flexible residues were modeled with continuous sidechain flexibility, and all other residues remained completely fixed.

The performance of desktop hardware appears similar to server hardware, except on a smaller scale. A single Nvidia GTX 1070 GPU performs minimizations at roughly half the speed of an Nvidia Tesla P100 GPU. Two Nvidia GTX 1080 GPUs perform similarly to the Nvidia Tesla P100 GPU on the large conformation benchmark (Fig. 3, bottom), but actually perform worse than a single Nvidia GTX 1070 for the small conformation benchmark (Fig. 3, middle) - despite having well over twice the hardware of the single Nvidia GTX 1070 GPU. This anomalous performance suggests the kernel OSPREY 3.0 uses for minimizations is not yet well-optimized for the Nvidia GTX 1080 GPU, and that future engineering efforts could offer significant performance increases for Nvidia GTX 1080 GPUs. The Nvidia GTX 1050, a laptop GPU, does not appear to be powerful enough to offer any advantages over traditional CPU computing in OSPREY 3.0 (Fig. 3, light blue columns).

Modern GPU architectures offer thousands of parallel hardware units for calculations, compared to the tens of parallel hardware units in modern CPU architectures. The performance results of the current generation of OSPREY’s GPU kernels indicate that minimization speeds on GPUs have only begun to scratch the surface of what is possible, particularly for minimizations with few atom pairs. Future versions of these GPU kernels will likely offer significantly higher performance on the same hardware - perhaps allowing minimization speeds many times faster than today’s GPU kernels. This in turn will make it even more efficient to perform minimization-aware protein design, and allow minimization-aware designs with even more mutable and flexible residues and with more mutation options per residue.

## PYTHON SCRIPTING IMPROVES EASE-OF-USE

One of the most visible additions to OSPREY 3.0 is the Python application programming interface (API), which allows fine-grained control over design parameters in a streamlined and easy-to-use experience. OSPREY 3.0 still supports a command-line interface with configuration files for backwards compatibility, but new development will be focused mostly on the new Python interface.

The OSPREY 3.0 distribution contains a Python module which is installed using the popular package manager pip. Once installed, using OSPREY 3.0 is as easy as writing a Python script. High-performance computations are still performed in the Java virtual machine to give the fastest runtimes, so Java is still required to run OSPREY 3.0, but communication between the Python environment and the Java environment is handled behind-the-scenes, and OSPREY 3.0 still looks and feels like a regular Python application.

See Figure 4 for a complete example of a Python script that performs a very simple design using OSPREY 3.0, and Figure 5 for a slightly more involved design using *BBK^*^*^36^ (a new algorithm in OSPREY 3.0, described in its own section below). Figure 6 graphically displays the design setup for the *BBK^*^* design.

**Figure 4:**
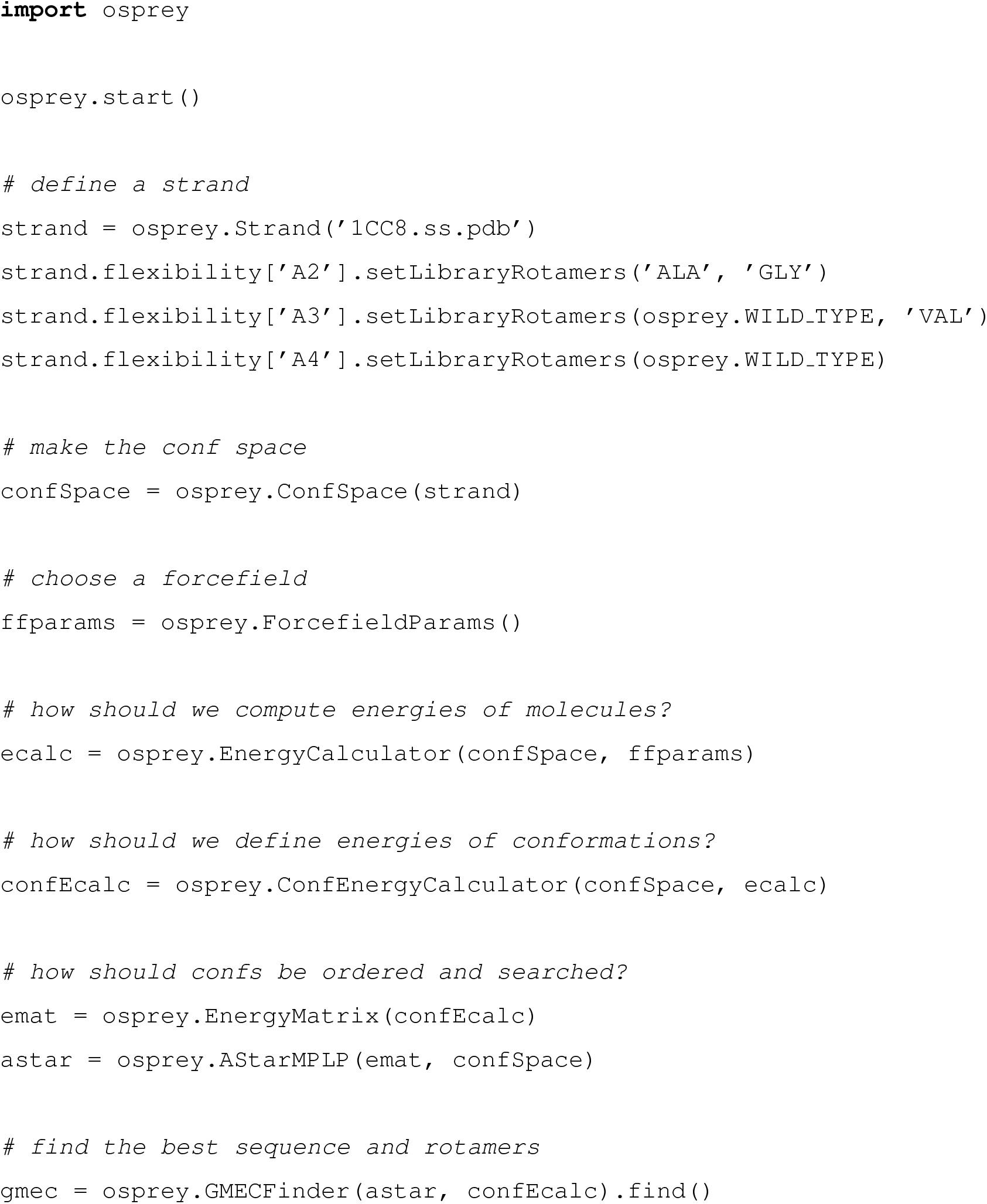
A Python script that performs a very simple design in OSPREY 3.0. The design searches over sequences in which residues A2 and/or A3 of the Atx1 metallochaperone protein (PDB ID: 1CC8)^47^ are mutated; residues A2-A4 (i.e., residue 2-4 of chain A) are all modeled with sidechain flexibility, consisting of a discrete search over the Penultimate rotamer library^48^’s rotamers for the specified amino acid types. The mutability, flexibility, and starting crystal structure are all specified in the “define a strand” section of the code. Advanced users can also modify the other sections to specify changes from the default search algorithms, energy function, and other modeling assumptions. This script uses the Max Product Linear Programming (MPLP) algorithm^49^ to reduce the size of the *A^*^* search tree^15^ employed for sequence and conformational search without compromising accuracy; see Ref. ^22^ for details.

**Figure 5:**
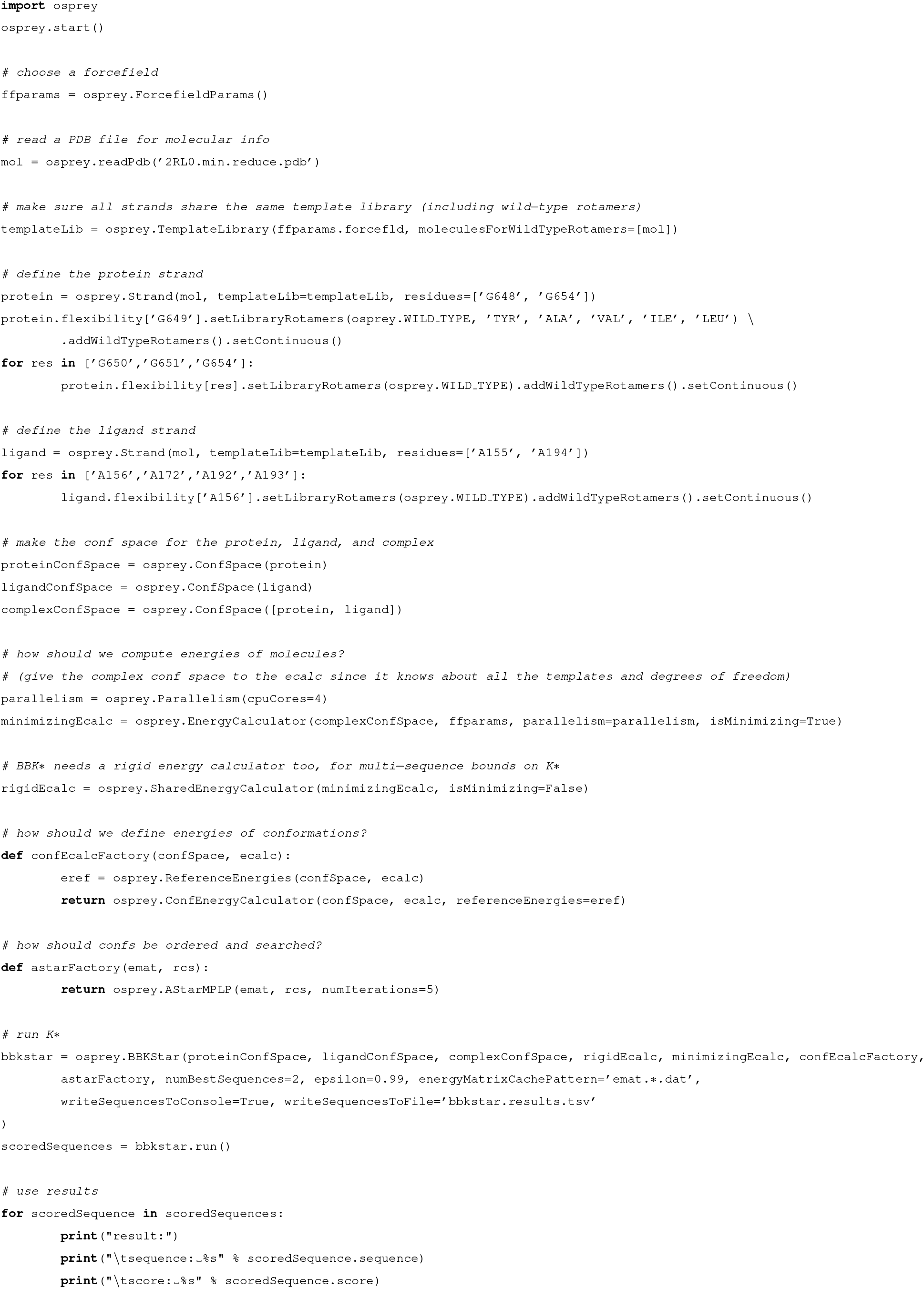
A Python script that performs a simple *BBK\** design in OSPREY 3.0. This design produces a peptide to bind human fibronectin (the “ligand strand,” i.e. chain A) by optimizing a fragment of the protein FnBPA from *Staphylococcus aureus* (the “protein strand,” chain G), which has been crystallized in complex with fibronectin domains (PDB ID: 2RL0^50^). As in Fig. 4, the script defines the starting crystal structure, mutable residues, and level of mutability and flexibility (here including continuous flexibility) in the form of Python strand objects. Fig. 6 represents this design graphically. This design is accelerated by parallelism, running on 4 CPU cores. This example thus shows it is easy to invoke and use parallelism within the OSPREY 3.0 software.

**Figure 6:**
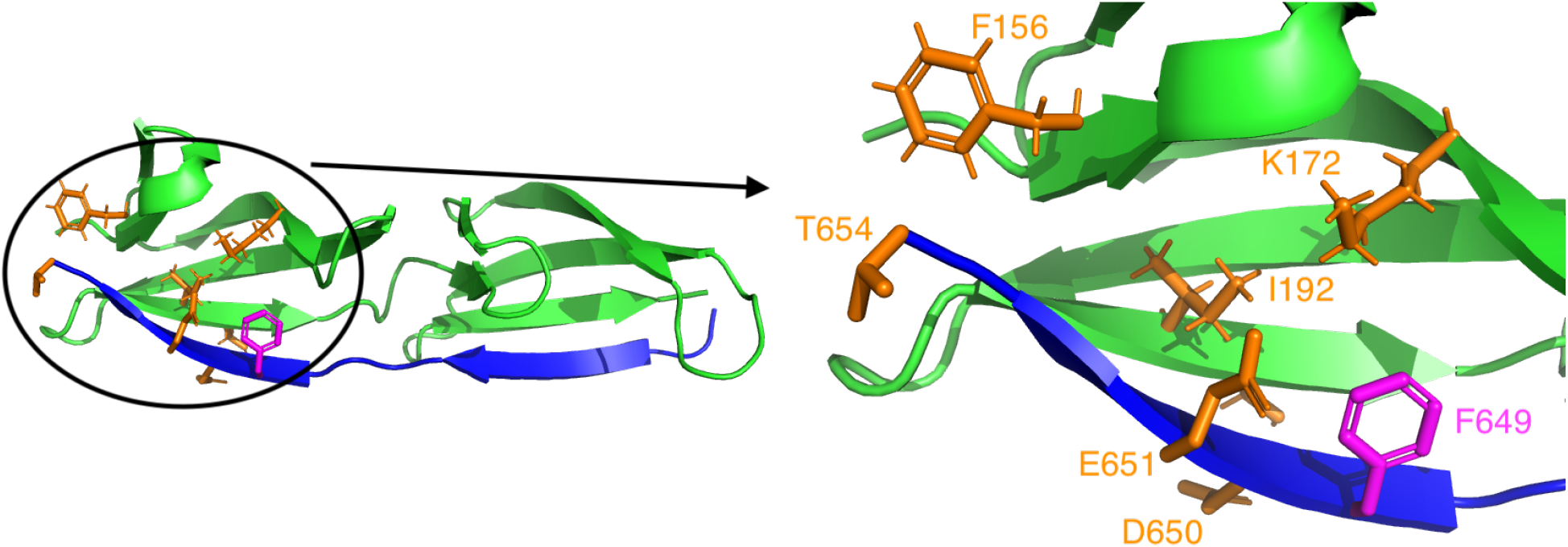
Setup for the Python-scripted *BBK*^*^^36^ design described in Fig. 5. This design starts with the crystal structure (PDB ID: 2RL0^50^) of a complex between fragments of the protein FnBPA from *Staphylococcus aureus* (blue ribbons) and human fibronectin (green ribbons), and optimizes binding with respect to the amino acid type of FnBPA residue 649 (magenta), while modeling continuous flexibility in several surrounding sidechains (orange). The full complex is shown on the left, while the region surrounding the mutation is shown in detail on the right. See Ref. 36 for background on the FnBPA:fibronectin system.

## NEW PROTEIN DESIGN ALGORITHMS IN osprey 3.0

### LUTE: Putting advanced modeling into a form suitable for efficient, discrete design calculations

OSPREY 3.0 comes with LUTE^18^, a new algorithm that addresses two issues with previous versions of OSPREY.

First, previous versions modeled continuous flexibility by enumerating conformations in order of a *lower bound* on minimized conformational energy^2,16^. This lower bound can be relative loose, especially for larger systems, and thus a large number of suboptimal conformations—often exponentially many with respect to the size of the system—must be scored by continuous minimization merely because they have favorable lower bounds on their energy. LUTE addresses this problem by enumerating conformations in order of their actual minimized conformational energies instead of simply in order of a lower bound. These energies are estimated using an expansion in low-order tuples of residue conformations. Thus, the burden of modeling continuous flexibility is shifted from the combinatorial optimization (*A^*^*) step, which has unfavorable asymptotic complexity, to a precomputation step (the “LUTE matrix precomputation” ^18^) that only scales quadratically with the number of residues. This dramatically reduces the computation time for large designs with continuous flexibility, and has doubled the number of residues that can be treated simultaneously with continuous flexibility^18^.

Second, all previous combinatorial protein design algorithms have relied on an explicit decomposition of the energy as a sum of local (e.g., pairwise) terms. This made design with energy functions that do not have this form difficult. LUTE can straightforwardly support general energy functions, and, as shown in Ref. 18, it can obtain good fits at least in the case of Poisson-Boltzmann energies. Moreover, once the LUTE matrix precomputation is completed, the time cost of finding the optimal sequence and conformation does not depend on the energy function used. This is an enormous advantage for more expensive and accurate energy functions like Poisson-Boltzmann, which otherwise would be far too expensive for all but the smallest designs.

OSPREY users can now turn on LUTE for continuously flexible calculations simply by setting a boolean flag (in the `DEEGMECFinder` Python constructor). OSPREY 3.0 also supports design with Poisson-Boltzmann solvation energy calculations, which call the DelPhi^51,52^ software for the single-point Poisson-Boltzmann calculations (we ask the user to download DelPhi separately for licensing reasons). Such improved modeling is essential to increasing the reliability of and range of feasible uses for computational protein design.

### CATS: Local backbone flexibility in all biophysically feasible dimensions

OSPREY pioneered protein design calculations that model local continuous flexibility of sidechains in the vicinity of rotamers in all biophysically feasible dimensions (i.e., the sidechain dihedrals). This continuous flexibility was often critical in correctly predicting energetically favorable sequences^1,16^, especially when those sequences falsely appeared to be sterically clashing when modeled using only rigid rotameric conformations taken from a rotamer library (see section on GPU acceleration above for more details). In OSPREY 3.0, we now extend this ability to the backbone: allowing local continuous backbone flexibility in the vicinity of the native backbone with respect to all biophysically feasible degrees of freedom.

This flexibility is enabled by the CATS algorithm^20^ (Fig. 7). CATS uses a new parameterization of backbone conformational space, along with the voxel framework that OSPREY has always included. It is equivalent to searching over all changes in backbone dihedrals (*ϕ* and *ψ*) subject to keeping the protein conformation constant outside of a specified flexible region. CATS includes an efficient Taylor series-based algorithm for computing atomic coordinates from its new degrees of freedom, enabling efficient energy minimization. Unlike previous protein design algorithms with backbone flexibility, CATS routinely finds backbone motions on the order of an angstrom (in RMSD with respect to the wildtype backbone) while still performing a comprehensive search of its backbone conformation space. In Ref. 20, we have shown that backbone flexibility as modeled by CATS is sometimes critical for avoiding nonphysical steric clashes (Fig. 7B,C) and often affects energetics significantly. For example, mutating residue 54 of the antibody VRC07 to tryptophan improves its binding to its antigen (HIV surface protein gp120)^7^, but a design to recapitulate this mutation found it to be blocked by a steric clash unless CATS was used to find a backbone motion that escapes the clash^20^. In this design, CATS significantly outperformed a provable search over backrub^53^ motions, which are also available in OSPREY^19,54^.

**Figure 7:**
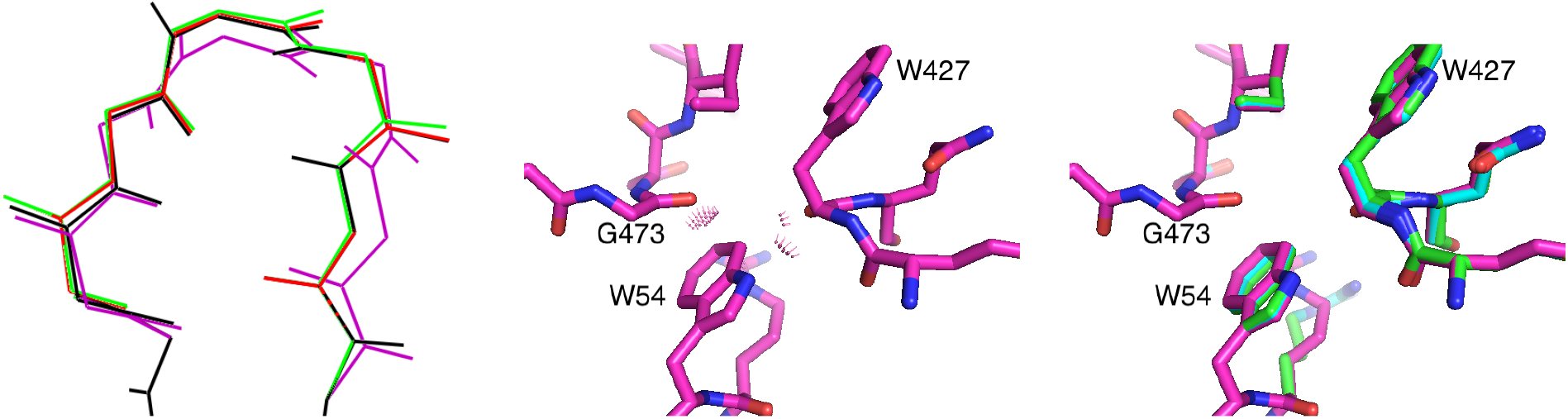
Left: CATS allows systematic search over a voxel of backbone conformations in the vicinity of the wild-type backbone conformation (black). The voxel is specified as box constraints on a novel set of backbone coordinates; conformations with one such coordinate moved to the edge of the voxel are shown in red and green, and a conformation with all such coordinates moved to the edge of the voxel is shown in purple. See Fig. 1 of Ref. 20 for more details. Middle: Rigid-backbone structural modeling of an experimentally effective mutant of anti-HIV gp120 antibody VRC07 showed unavoidable steric clashes between Trp 54 of VRC07 and Trp 427 and Gly 473 of gp120 (purple). Right: CATS explained the experimentally observed activity by finding a new backbone conformation that resolved these clashes (green; overlaid with clashing rigid-body backbone (purple) and backbone conformation computed with the older DEEPer algorithm (blue)). DEEPer reduced the clashes somewhat using backrub motions^53^, but they were still significant even after the backrubs. See Fig. 3 of Ref. 20 for more details. Portions of this figure were reprinted with permission from Ref. 20.

CATS is intended to be run as part of the flexibility model for OSPREY’s other algorithms, yielding efficient calculations with continuous flexibility in both the sidechains and the backbone. OSPREY’s convenient interface allows a user to add CATS flexibility to a design merely by specifying the start and end points of the backbone segment to be made flexible.

### BBK^*^: Efficiently computing the tightest binding sequences from a combinatorially large number of binding partners

In previous versions of OSPREY, the *K^*^* algorithm^24^ modeled an ensemble of Boltzmann-weighted conformations to approximate the thermodynamic partition function. It combined minimized dead-end elimination pruning^14^ with *A^*^* ^14,55^ gap-free conformation enumeration to compute provable *ε*-approximations to the partition functions for the protein and ligand states of interest. *K^*^* combined these partition function scores to approximate the association constant, *K_a_*, as the ratio of *ε*-approximate partition functions between the bound and unbound states of a protein-ligand complex. Notably, each partition function ratio, called a *K^*^ score*, is provably accurate with respect to the biophysical *input model*^2,16,24^.

Although *K^*^* efficiently and provably approximated *K_a_* for a given sequence, it had to compute a *K^*^* score for each sequence of interest. All provable ensemble-based algorithms prior to *BBK^*^*, as well as many heuristic algorithms that optimize binding affinity, are *single-sequence* algorithms which must compute the binding affinity for each possible sequence. The number of sequences, of course, is exponential in the number of simultaneously mutable residue positions. Therefore, designs with many mutable residues rapidly became intractable. OSPREY 3.0 provides a new algorithm, *BBK^*^*, which overcomes this challenge. *BBK^*^*^36^ builds on *K^*^*, and is the first provable, ensemble-based protein design algorithm to run in time sublinear in the number of sequences. The key innovation in *BBK\** that enables this improvement is the *multi-sequence (MS) bound.* Rather than compute binding affinity separately for each possible sequence, as single-sequence methods do, *BBK^*^* efficiently computes a single provable *K^*^* score upper bound for a combinatorial number of sequences. *BBK\** uses MS bounds to prune a combinatorial number of sequences during the search, entirely avoiding single-sequence computation for all pruned sequences.

Importantly, *BBK^*^* also contains many other powerful algorithmic improvements and implementation optimizations: the parallel architecture of *BBK\**, which enables concurrent energy minimization, and a novel two-pass partition function bound, which minimizes far fewer conformations while still computing a provable *ε*-approximation to the partition function. Combined with the combinatorial pruning power of the MS bound, *BBK\** is able to search over much larger sequence spaces than previously possible with single-sequence *K^*^* (Fig. 8). In computational experiments on 204 protein design problems, *BBK^*^* accurately predicted the tightest-binding sequences while only computing *K^*^* scores for as few as one in 10^5^ of the sequences in the search space^36^. Moreover, in computational experiments on 51 protein-ligand design problems, *BBK\** was up to 1982-fold faster than single-sequence *K^*^*, despite provably producing the same results ^36^.

**Figure 8:**
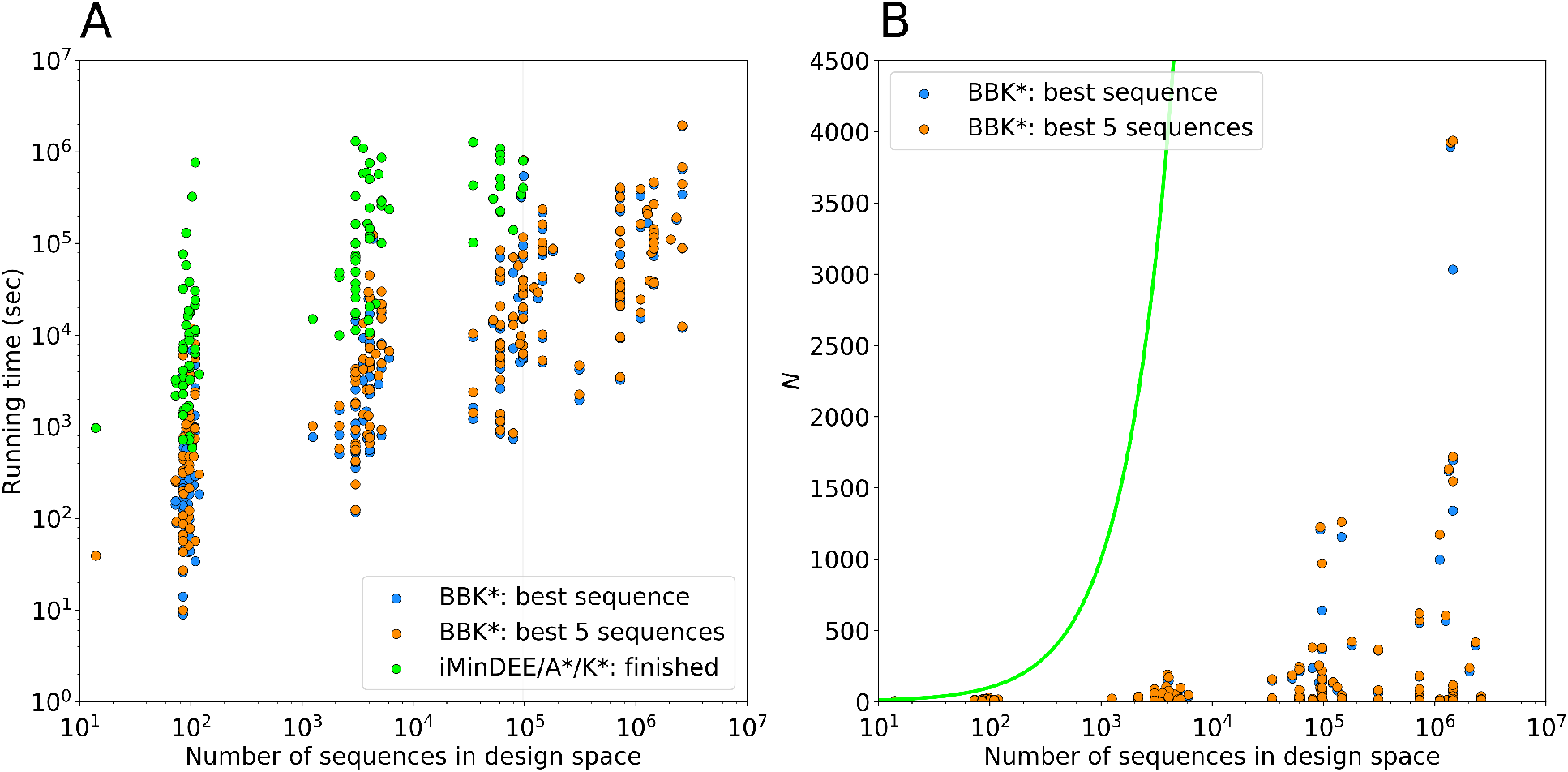
Measuring the contributions of the *BBK^*^*^36^ algorithmic improvements to the empirically observed running times of OSPREY. *BBK^*^* calculations were run to predict either the single top sequence or to enumerate the top 5, and were compared to exhaustive computation of *K^*^* scores for each sequence (i.e., iMinDEE/*A^*^*/*K^*^*^2,16^ or singlesequence *K^*^*), which was the prior state of the art for Boltzmann-weighted ensemble-based binding affinity computation before *BBK*\*. (A) Running times for *BBK^*^* and single-sequence *K^*^* vs. the number of sequences in the search space for 204 protein design test cases, a benchmark set described in Ref. 36. Single-sequence *K^*^* completed only 107 of the test cases within a 30-day time limit (left of the vertical line), and took up to 800 times longer than *BBK^*^* to do so, while *BBK^*^* completed all the designs within the time limit. (B) The number N of sequences whose energies must be examined or bounded by iMinDEE/*A^*^*/*K^*^* (green line; exponential in the number of mutable residue positions) and by *BBK^*^* (dots). For each data point representing a *BBK^*^* test case, the vertical gap between that data point and the green line (gap on the *y* axis) represents the number of sequences that are pruned without ever having to be examined. Figure adapted with permission from Ref. 36.

These improvements show that *BBK^*^* not only accelerates protein designs that were possible with previous provable algorithms, it also efficiently performs designs that are too large for previous methods.

### BWM^*^: Exploiting locality of protein energetics to efficiently compute the GMEC

OSPREY 3.0 comes with BWM^*^^23^, a new algorithm that exploits sparse energy functions to provably compute the GMEC in time exponential in merely the branch-width w of a protein design problem’s sparse residue interaction graph.

Because energy decreases as a function of distance, many protein design algorithms model protein energetics with energy functions which omit pairwise interactions between sufficiently distant residues. These *sparse energy functions* not only provide a simpler, more efficiently computed model of energy, but also induce *optimal substructure* to the problem: because not all residues interact, the optimal conformation for a given residue can be independent of the conformations at other residues. BWM^*^ exploits this optimal substructure by 1) representing the sparse interactions with a sparse residue interaction graph, and 2) computing a branch-decomposition for use in dynamic programming.

BWM^*^, unlike treewidth-based methods that also exploit the sparsity of pairwise residue interactions to efficiently compute the GMEC ^56^, enumerates a gap-free list of conformations in order of increasing sparse energy. Because this list is gap-free, BWM^*^ not only computes the GMEC of the sparse energy function, but also recovers the GMEC of the full energy function, as shown in Ref. 23. By enumerating all conformations within the provable sparse energy bound between the sparse and full GMEC, BWM^*^ computes a list of conformations that is guaranteed to contain the full GMEC, as well as the sparse GMEC^57^. Moreover, because BWM^*^ can enumerate conformations in gap-free order up to any energy threshold specified by the user, it can be used to accurately compute partition functions, and thus binding free energies that account correctly for entropy, using the *K^*^* algorithm^2,24^.

Thus, in practice, BWM^*^ circumvents the worst-case complexity of traditional methods such as *A\** for designs with sparse energy functions, computing the sparse GMEC of an *n*-residue design with at most *q* rotamers per residue in 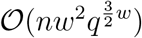 time, and also enumerates each additional conformation in merely 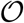(*n* log *q*) time, which is up to three orders of magnitude faster than traditional *A^*^* in practice^23^.

## ACCURACY BENCHMARKS

We first tested the accuracy of OSPREY 3.0 for the subset of algorithms also available in OSPREY 2.2,*β*, by running both versions of OSPREY on the same test cases and checking that the results matched. Since the accuracy of OSPREY 2.2*β* using these algorithms has been experimentally confirmed (see Introduction), by transitivity, our tests confirmed OSPREY 3.0’s accuracy. In addition, we performed new, retrospective tests, described below.

To evaluate the accuracy of the implementation of the newest optimizations in OSPREY 3.0, we performed a series of designs for a variety of protein-protein interfaces (PPIs) as retrospective validation. We used *K*\*^24^ to computationally predict experimentally measured changes in binding for each PPI. Each protein structure is listed by name and PDB ID in Table 2^58–61^. These systems include barnase with its peptide inhibitor barstar^62,63^, the cytochrome *c*:cytochrome *c* peroxidase complex^64^, interferon *α*-2 (IFN*α*2) in complex with interferon *α/β* receptor 2 (IFNAR2)^65^, and the interleukin 2 (IL-2):IL-2 receptor *α* (IL-2R*α*) complex ^66^.

**Table 2:**
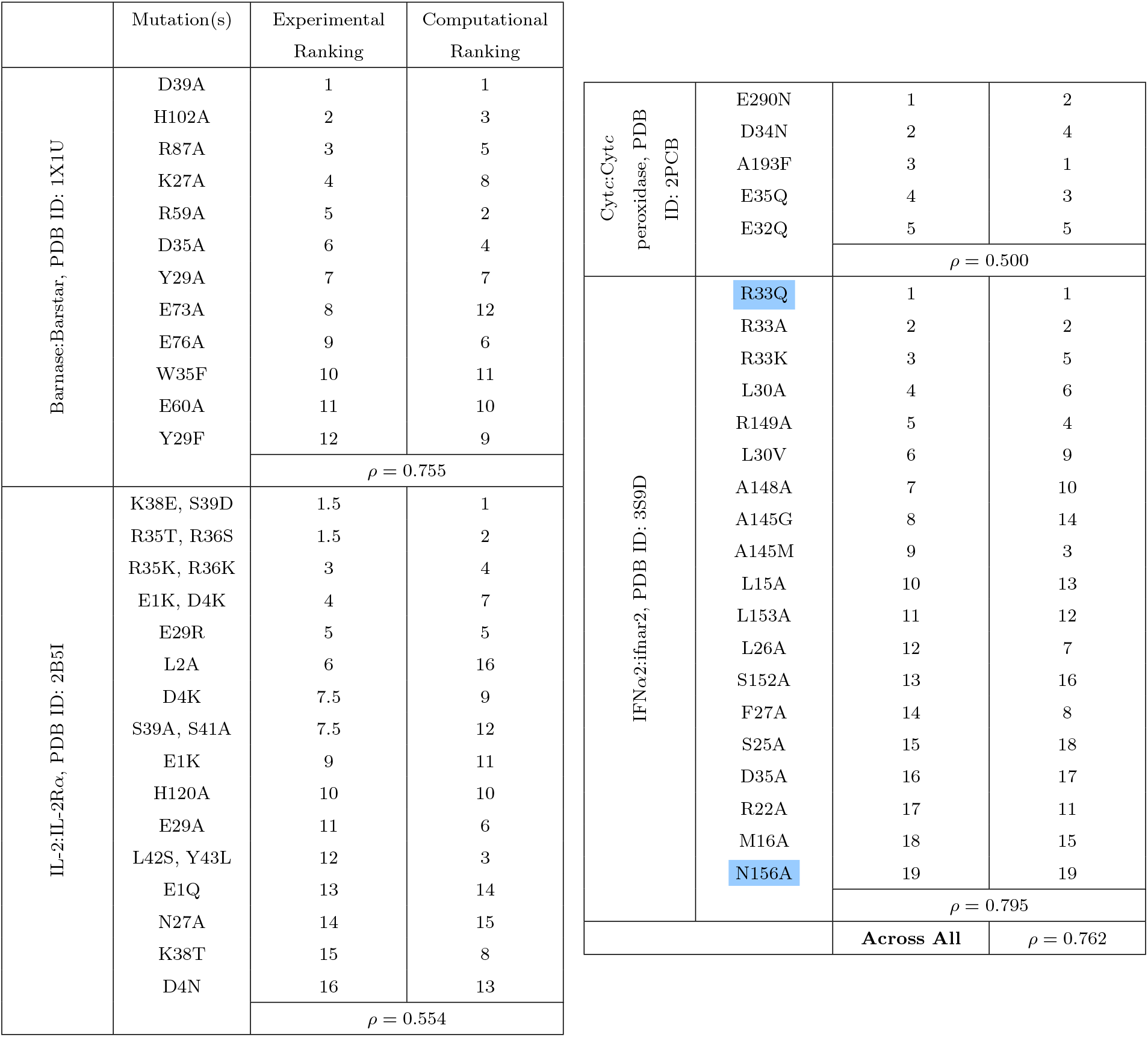
Comparison of OSPREY predictions to experimental results for mutations in four protein systems. Allowed mutations for each system are listed along with their corresponding rankings from experimental measurements^62–66^ vs. computational predictions by *K^*^* from OSPREY 3.0. The mutations highlighted in blue are shown in detail in Figure 9. A Spearman’s *ρ* value is calculated for each system and shown here. The “Across All” value is calculated by ranking each system individually and then calculating the Spearman’s *ρ* across all of the designs.

Our retrospective validation experiments focused on mutations at residues in or proximal to the protein-protein interface that were not limited to alanine scanning. Including some of these tested and reported mutations^62–66^, for each structure we tested anywhere from 5 to 19 designs. In total, we tested 58 mutations using default, out-of-the-box OSPREY 3.0 settings and parameters. Each design included one or two mutable residues along with a set of surrounding flexible residues (See Table 2). Flexible residues were chosen by selecting all residues within 4 Ǻ of the mutable residues and removing those that only have backbone interactions. Two example designs are shown in Figure 9, where OSPREY 3.0 and *K*^*^ accurately predict the effect of two point mutations in the interface of the IFN*α*2:IFNAR2 complex (highlighted in blue in Table 2).

**Figure 9:**
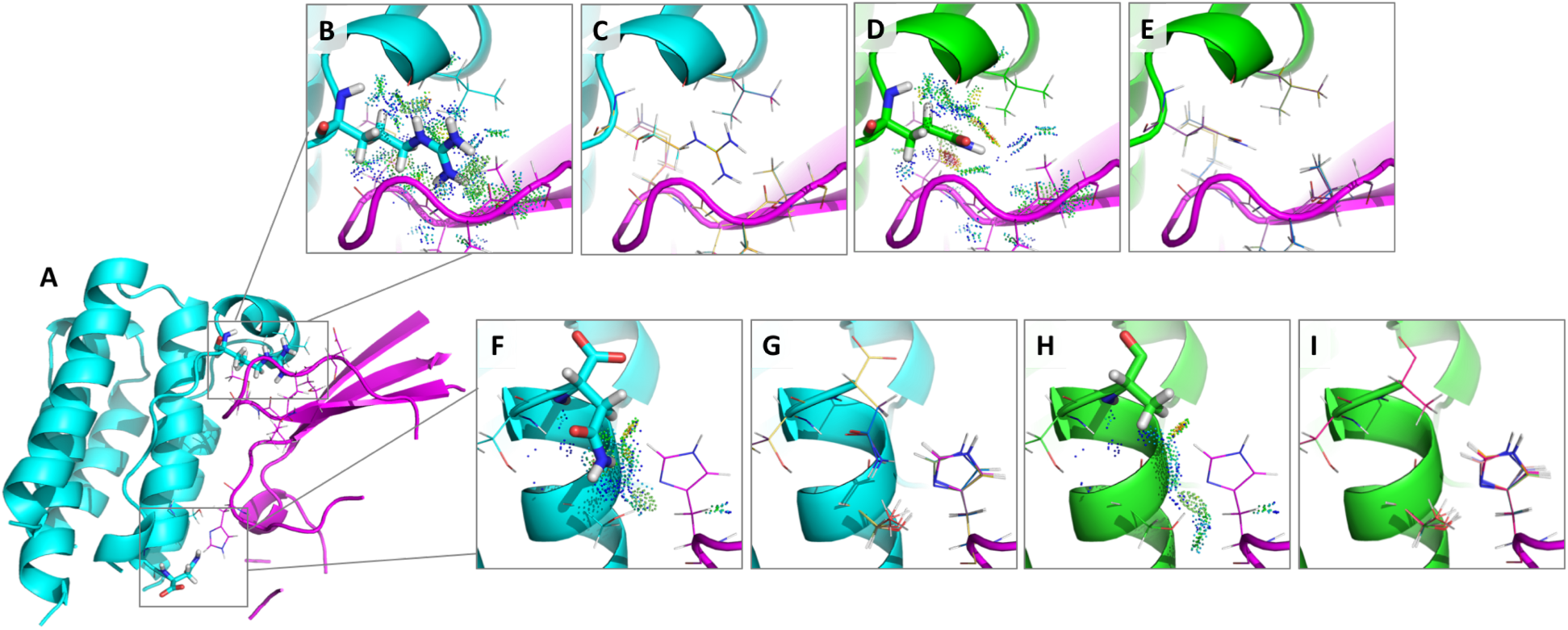
(A) The structure of the IFNa2:IFNαR2 complex (PDB ID: 3S9D ^60^) with separate chains shown in cyan and magenta and with two example interface design regions shown in boxes. Each box contains a mutable residue shown as sticks and its surrounding flexible residues shown as lines. (B) and (F) zoom in on each design. (B-E) Design at position R33 for a mutation that OSPREY correctly predicts as decreasing binding: R33Q. (B) The wildtype sequence with probe dots^67,68^ displaying favorable interactions with surrounding flexible residues (shown as lines). (D) The mutant sequence (33Q) with probe dots displaying some favorable as well as unfavorable interactions. Comparing (B) and (D), it is clear there is a loss in favorable interactions and a gain in unfavorable interactions upon mutation from R to Q, resulting in an experimentally observed decrease in binding that the *K^*^* algorithm captures accurately (See Table 2). (C) and (E) show the top 10 conformations in the conformational ensemble used in the *K^*^* calculation for each sequence. (F-I) Design at position N156 for a mutation that OSPREY correctly predicts as increasing binding: N156A. (F) The wildtype sequence with probe dots ^67,68^ displaying some favorable interactions with surrounding flexible residues (shown as lines). (H) The mutant sequence (156A) with probe dots displaying some favorable interactions with surrounding flexible residues (shown as lines). There are some gained interactions (shown by an increase in the number of favorable probe dots) in (H) compared to (F), but these are not visually obvious, thus emphasizing the importance of *K^*^*, which successfully picks up these nuanced changes and correctly predicts improved binding (See Table 2). (G) and (I) show the top 10 conformations in the conformational ensemble used in the *K^*^* calculation for each sequence. Not shown are the ensembles for the unbound states that are also used to calculate the *K^*^* scores.

For each system, the *K^*^* scores were ranked in increasing order of reported experimental binding. Spearman’s *ρ* values were subsequently calculated for each system by calculating the statistical dependence between the *K^*^* score rankings and the experimentally measured rankings (See Table 2 and Figure 10). This is a sound measure because generally the output of a design calculation that is used to decide which mutants to make experimentally is simply the intra-system ranks of the mutants. Looking at the values in Table 2, we see a high correlation in the rankings between experimentally measured binding and binding predicted by OSPREY 3.0 and *K^*^* for each system with values ranging from 0.500 to 0.795. We found that, across the tested systems, the Spearman’s *ρ* value is 0.762. This value is the Pearson correlation of the intra-system ranks of all the mutants. Overall, these correlations are very good for design for affinity in computational protein design.

**Figure 10:**
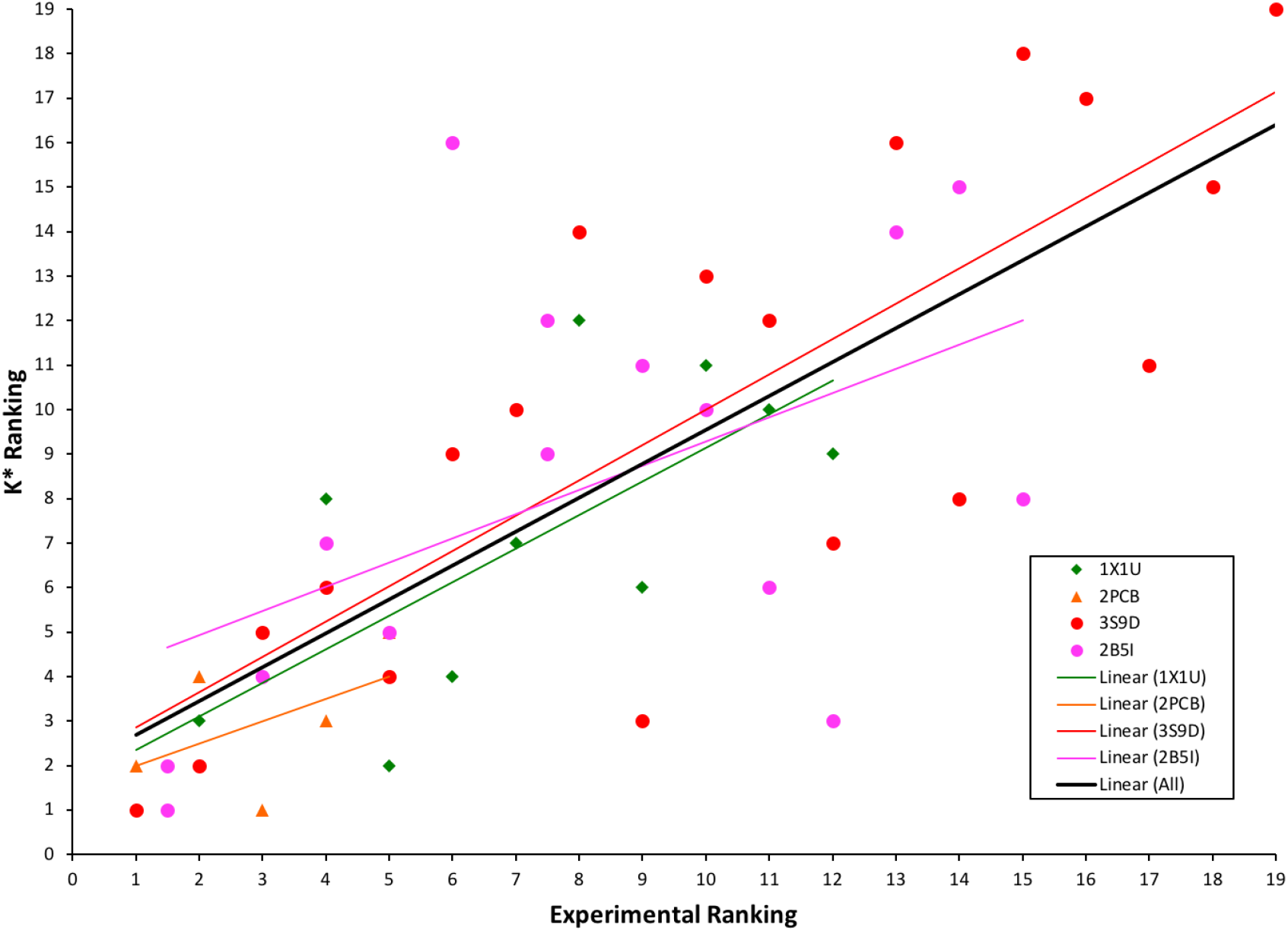
Testing the accuracy of the *K^*^* algorithm in OSPREY 3.0 by comparing *K^*^* rankings to experimentally reported rankings (See Table 2). Each system is represented by its corresponding PDB ID and a linear trendline is shown for each in its corresponding color according to the legend.

## DISCUSSION

OSPREY has demonstrated its accuracy and utility in practice through many prospective designs that have performed well experimentally^6–12^. OSPREY 3.0 is at least as accurate as the versions of OSPREY used to perform these designs, because it uses the same biophysical model used in those studies, with provable guarantees of accuracy given the biophysical model. We have compared design results using OSPREY 2.2 and OSPREY 3.0 to confirm agreement. However, OSPREY 3.0 performs such designs much more efficiently, due to the engineering improvements described here. Moreover, in this paper we have performed additional comparisons to experimental data to confirm the accuracy of OSPREY 3.0. OSPREY 3.0 also includes methods to improve the biophysical model and thus improve accuracy still further (should the user choose to select OSPREY’s newer models).

As our benchmark results here show, we have made substantial progress toward correctly predicting the effect of mutations on protein activity. The high accuracy comes from OSPREY’s accurate biophysical model, which accounts for both continuous protein flexibility and conformational entropy, together with algorithms that provably return optimal sequences given that model. In fact, no other software can provide a provable guarantee of accuracy given a model that accounts for continuous flexibility and conformational entropy. Moreover, OSPREY’s combinatorial algorithms^4,5^ compute optimal sequences efficiently even when searching over a large sequence space.

The large speedups in OSPREY 3.0, together with the easy-to-use Python interface, thus make it much more tractable to perform protein design with such biophysically realistic modeling and with guaranteed accuracy given the model. In particular, OSPREY 3.0 benefits from many sources of speedups that can be used together. Speedups from OSPREY 3.0’s optimization of the conformational minimization, forcefield evaluation, and A* routines can exceed two orders of magnitude even compared to OSPREY 2.2^21^ running on the same CPU hardware. Together with an additional speedup of over an order of magnitude from GPU’s, a design that would take months using OSPREY 2.2 could easily take only a few hours using OSPREY 3.0. Many designs could see even greater speedups, because in addition to these engineering improvements, some of the algorithmic improvements in OSPREY 3.0 provide a dramatic increase in computational efficiency.

The improvements in modeling facilitated by OSPREY 3.0’s new algorithms also make protein design with OSPREY more realistic. However, there is still much room for improvement in the biophysical model used by OSPREY, and indeed by all currently available protein design software. Modeling of larger backbone motions, more realistic interactions with water, and electronic polarization, among other phenomena, are all likely to yield substantial improvements in accuracy. The refactored architecture of OSPREY 3.0 will make it easier to experiment with algorithms that facilitate these modeling improvements, and to implement these algorithms within OSPREY’s current code base. Moreover, we have released OSPREY 3.0 as open source, to aid the community both in the development and the application of improved models and algorithms for computational protein design.

## CONCLUSIONS

OSPREY has long offered unique capabilities to protein designers. In particular, it has always offered a unique combination of provably accurate conformational search, continuous flexibility, efficient search over large sequence spaces, and free energy calculations based on Boltzmann-weighted thermodynamic conformational ensembles. In OSPREY 3.0 we introduced software improvements that will make these algorithms much more practical for the wider design community: performance that is orders of magnitude faster, and a Python interface that makes OSPREY much easier to use. In addition, we expanded the range of biophysical modeling assumptions that OSPREY can accommodate, both in terms of molecular flexibility and energy functions. As with previous versions, we are releasing OSPREY 3.0 as free and open-source software to maximize its benefit to the community. We hope this new version will be of significant utility to designers, whether they have used OSPREY before or are trying it for the first time.

## ACKNOWLEDGMENTS

The authors would like to thank Dr. Alvin Lebeck for helpful discussions on GPUs, Drs. Kyle Roberts and Swati Jain for helpful discussions on protein design, and the NIH (grants R01 GM-78031 and R01 GM-118543 to B.R.D.), NSF (Graduate Research Fellowship to A.O.), PhRMA Foundation (Informatics Predoctoral Fellowships to A.U.L. and M.A.H.), and Liebmann Foundation (fellowship to M.A.H.) for funding.

